## Supplement for "Integration of parallel pathways for flight control in a hawkmoth reflects prevalence and relevance of natural visual cues"

**Supplementary Figures**

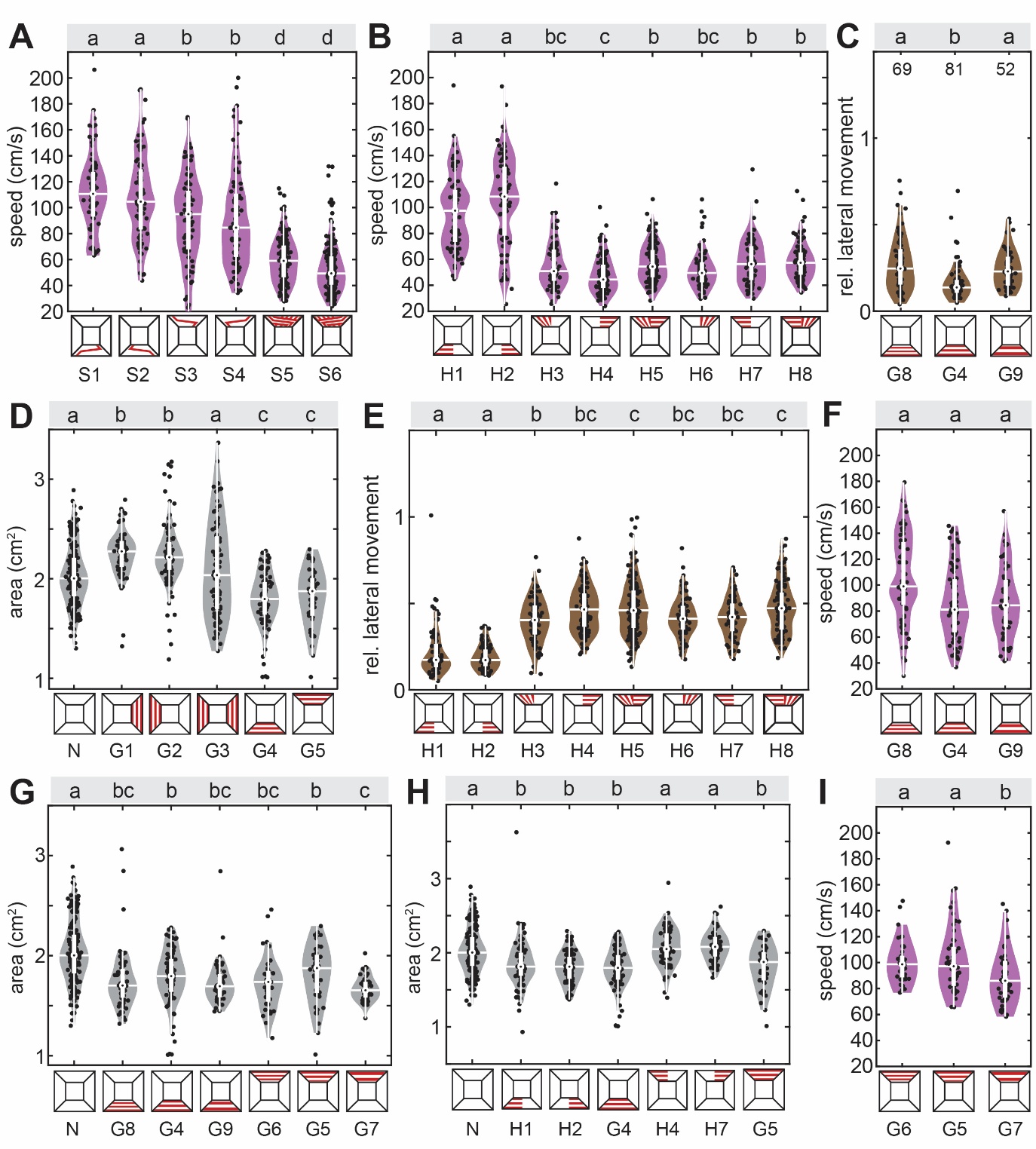

Fig. S1 **Optic flow based flight control and dorsal directional responses.**

**A** Average speed of flight paths with a red stripe which changed its position in the central third of the tunnel, crossing from one tunnel side to the other. The last two conditions present a version of this stripe, which repeated at the same frequency as the grating patterns. **B,E** Average speed and proportion of lateral movement of flight paths with gratings perpendicular (generating strong translational optic flow) and parallel (weak translational optic flow) to the tunnel’s longitudinal axis, covering one side of either the tunnel ceiling or floor. **C, F, I** Proportion of lateral movement and average speed with gratings of different spatial frequencies (repeating every 3 cm, 6 cm and 12 cm), mounted on the tunnel floor or ceiling, respectively. **D, G, H** Area of hawkmoth silhouette in tunnel videos, as a measure of their flight height above the tunnel floor (the smaller the area, the higher the hawkmoths) with grating patterns **D** on either tunnel side, **G** gratings of various spatial frequencies mounted ventrally and dorsally, and **H** gratings covering half the tunnel or the full tunnel ventrally and dorsally. Black letters show statistically significant differences in group medians (confidence level: 5%). Conditions with different letters were significantly different from each other. The white boxplots depict the median and 25% to 75% range, the whiskers represent the data exceeding the box by more than 1.5 interquartile ranges and the violin plots indicate the distribution of the individual data points shown in black. For the letter-number identifiers below each condition see table S1.

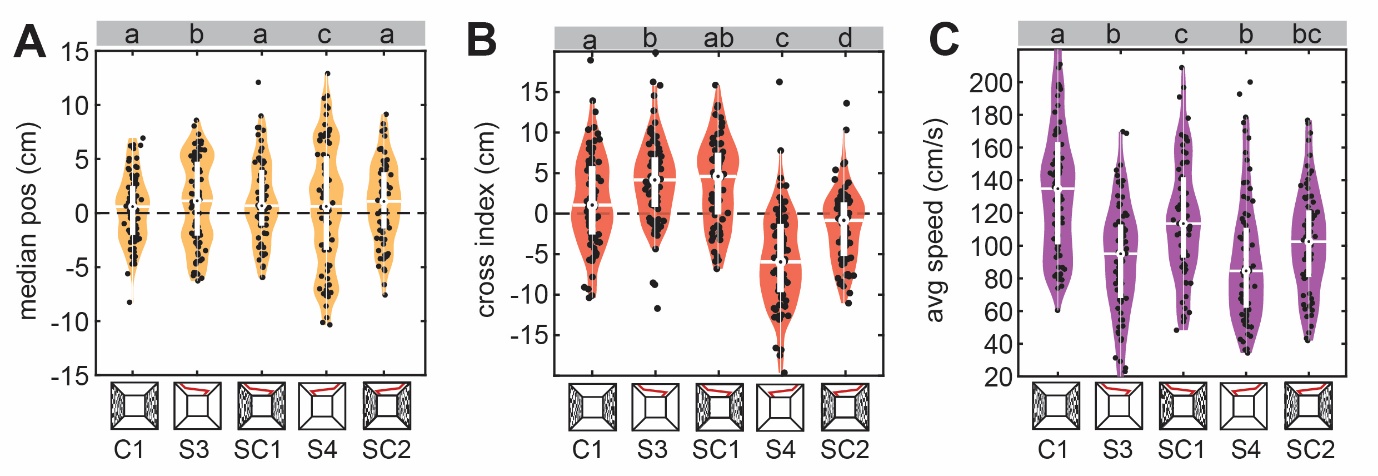

**Fig. S2** **Cue conflict: lateral optic flow and dorsal directional cues.**

**A-C** Median lateral position, cross-index and average speed of flight paths with a red stripe which changed its position in the central third of the tunnel, crossing from one tunnel side to the other, and a 50% contrast checkerboard pattern on both lateral tunnel sides, presented individually and in combination. Black letters show statistically significant differences in group medians (confidence level: 5%). Conditions with different letters were significantly different from each other. The white boxplots depict the median and 25% to 75% range, the whiskers represent the data exceeding the box by more than 1.5 interquartile ranges and the violin plots indicate the distribution of the individual data points shown in black.

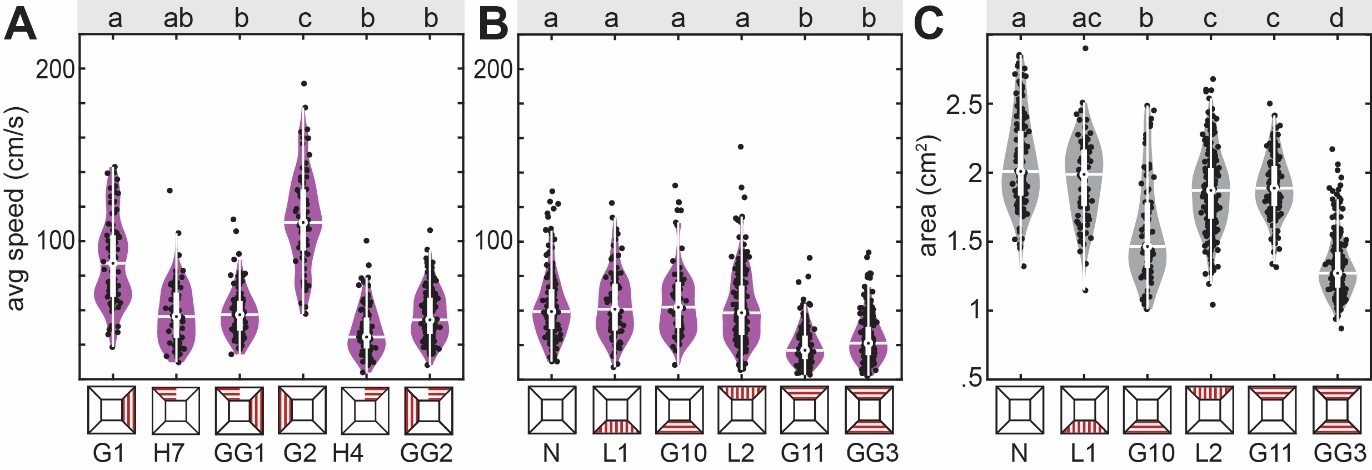

**Fig. S3** **Cue conflict: lateral distance regulation versus dorsal avoidance. A** Average speed of flight paths with lateral gratings and dorsal gratings covering half the tunnel, presented individually and in combination, and for **B** dorsal and ventral gratings perpendicular (optic-flow inducing) and parallel to the flight direction. **C** Average area of hawkmoths as a readout for flight height (the smaller the area, the higher the hawkmoths in the tunnel) in videos with dorsal and ventral longitudinal and perpendicular gratings. Black letters show statistically significant differences in group medians (confidence level: 5%). Conditions with different letters were significantly different from each other. The white boxplots depict the median and 25% to 75% range, the whiskers represent the data exceeding the box by more than 1.5 interquartile ranges and the violin plots indicate the distribution of the individual data points shown in black.

**Supplementary Tables**

**Table S1** Summary of visual stimulation conditions (labels as used in data repository <https://figshare.com/s/e680da3be83fe172a5e4>), and number of flight tracks per condition.

|  | **Condition short label** | **Condition label in  the repository** | **Condition description** | **flight track number (N)** |
| --- | --- | --- | --- | --- |
| 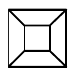 | N | none | no optic flow or contrast cues | 160 |
| 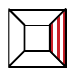 | G1 | right OF | right optic flow | 81 |
| 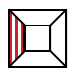 | G2 | left OF | left optic flow | 79 |
| 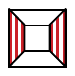 | G3 | lateral OF | lateral optic flow | 80 |
| 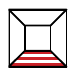 | G4 | ventral OF | ventral optic flow | 81 |
| 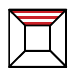 | G5 | dorsal OF | dorsal optic flow | 82 |
| 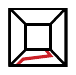 | S1 | ventral switch lr | ventral stripe, switch from left to right | 80 |
| 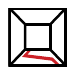 | S2 | ventral switch rl | Ventral stripe, switch from right to left | 81 |
| 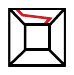 | S3 | dorsal switch lr | dorsal stripe, switch from left to right | 80 |
| 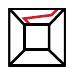 | S4 | dorsal switch rl | dorsal stripe, switch from right to left | 80 |
| 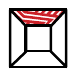 | S5 | dorsal switchgrating lr | dorsal switchgrating from left to right | 174 |
| 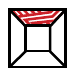 | S6 | dorsal switchgrating lr | dorsal switchgrating from right to left | 196 |
| 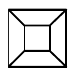 | N | none2 | none2 | 156 |
| 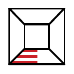 | H1 | ventral half OF l | ventral half optic flow left | 81 |
| 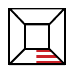 | H2 | ventral half OF r | ventral half optic flow right | 82 |
| 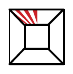 | H3 | dorsal half long l | dorsal half parallel grating left | 87 |
| 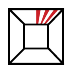 | H6 | dorsal half long r | dorsal half parallel grating right | 82 |
| 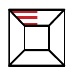 | H7 | dorsal half OF l | dorsal half optic flow left | 81 |
| 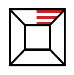 | H4 | dorsal half OF r | dorsal half optic flow right | 88 |
| 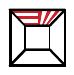 | H8 | dorsal half OF l long r | dorsal half optic flow left parallel grating right | 203 |
| 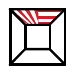 | H5 | dorsal half long l OF r | dorsal half parallel grating left optic flow right | 126 |
| 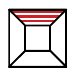 | G6 | dorsal OF 3cm | dorsal optic flow 3cm period | 51 |
| 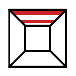 | G7 | dorsal OF 12cm | dorsal optic flow 12cm period | 50 |
| 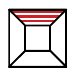 | G8 | ventral OF 3cm | ventral optic flow 3cm period | 69 |
| 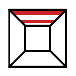 | G9 | ventral OF 12cm | ventral optic flow 12cm period | 52 |
| 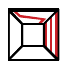 | SG1 | dorsal switch lr right OF | dorsal stripe, switch from left to right and right optic flow | 80 |
| 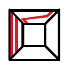 | SG2 | dorsal switch rl left OF | dorsal stripe, switch right to left and left optic flow | 79 |
| 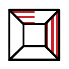 | GG1 | dorsal half OF l right OF | dorsal half optic flow left and lateral optic flow right | 126 |
| 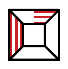 | GG2 | dorsal half OF r left OF | dorsal half optic flow right and  lateral optic flow left | 203 |
| 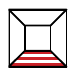 | G10 | ventralOF2 | ventral optic flow 2 | 69 |
| 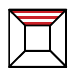 | G11 | dorsalOF2 | dorsal optic flow 2 | 157 |
| 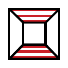 | GG3 | dorsal OF ventral OF | dorsal optic flow and ventral optic flow | 227 |
| 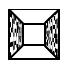 | C1 | lateral checker | lateral random checker | 81 |
| 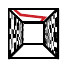 | SC1 | lateral checker switch lr | lateral random checker and dorsal stripe, switch from left to right | 73 |
| 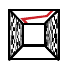 | SC2 | lateral checker switch rl | lateral random checker and dorsal stripe, switch from right to left | 67 |
| 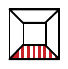 | L1 | ventral long | ventral parallel grating | 69 |
| 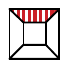 | L2 | dorsal long | dorsal parallel grating | 235 |
